# Long-term changes in kelp forests in an inner basin of the Salish Sea

**DOI:** 10.1101/2020.02.13.947309

**Authors:** Helen D. Berry, Thomas F. Mumford, Bart Christiaen, Pete Dowty, Max Calloway, Lisa Ferrier, Eric E. Grossman, Nathan R. VanArendonk

## Abstract

Understanding the historical extent of biogenic habitats can provide insight into the nature of human impacts and inform restoration and conservation actions. Kelp forests form an important biogenic habitat that responds to natural and human drivers. Global concerns exist about threats to kelp forests, yet long term information is limited and research suggests that trends are geographically distinct. We examined distribution of the bull kelp *Nereocystis luetkeana* over 145 years in South Puget Sound (SPS), a semi-protected inner basin in a fjord estuary complex in the northeast Pacific Ocean. We synthesized 48 historical and modern *Nereocystis* surveys and examined presence/absence within 1-km shoreline segments along 452 km of shoreline. Over the last 145 years, *Nereocystis* has been documented in 26% of the shoreline segments. Its extent decreased 62% basin-wide between the 1870s and 2017, with extreme losses in the two out of three sub-basins (96% in Central and 83% in West). In recent years, almost all *Nereocystis* occurred in the East sub-basin. In the majority of segments where *Nereocystis* disappeared, the most recent observation was 4 decades ago, or earlier. Multiple natural and human factors that are known to impact kelp could have contributed to observed patterns, but limited data exist at the spatial and temporal scale of this study. In some areas, recent environmental conditions approached thresholds associated with decreased kelp performance. Longstanding *Nereocystis* losses occurred exclusively in areas with relatively low current velocities. Remaining *Nereocystis* predominantly occurred in areas where circulation is stronger. Exceptions to this pattern demonstrate that additional factors outside the scope of this study contributed to trajectories of *Nereocystis* persistence or loss.

## Introduction

Humans have altered coastal ecosystems for centuries, yet we frequently lack long-term datasets to define a baseline that precedes significant human impacts and to identify changes from that baseline. The need for long-term reference points was initially identified in the context of global fisheries [1], but it is equally important to understand changes in biogenic habitats as losses can trigger changes to broader ecosystem structure and services. Historical maps have been used to estimate changes in the spatial extent of coastal habitats such as salt marshes, oyster reefs, coral reefs and kelp forests over century time scales [2–5]. Extended temporal baselines can enhance understanding of habitat variability and change. For example, historical nautical charts were used to spatially and temporally extend baseline information for coral reef extent in the Florida Keys, revealing greater loss than was previously quantified, along with newly identified areas of intact offshore reefs [4]. Enhanced understanding of the magnitude and spatial patterns of change in biogenic habitats can guide restoration and conservation actions, inform research into stressors, and support evaluation of changes to species that rely on these habitats.

Kelp forests occur predominantly in temperate oceans. They provide habitat for a wide range of species, especially invertebrate and fish assemblages [6]. Kelp forests are considered ecosystem engineers [7] because they create structural habitat with distinct local conditions by modifying the physical environment, such as light, water flow, sedimentation and pH [8, 9]. Extremely high productivity rates create habitat and food for local and distant food webs [10, 11]. Because kelp generally requires cold and nutrient-rich water [12, 13], large-scale climate cycles or changes can influence kelp abundance [5, 14, 15]. Grazing from herbivores also strongly influences kelp distribution and abundance[16], especially when changes in predator populations trigger linked changes in grazers[6, 17, 18].

Recent kelp forest losses across the globe have generated widespread concern. A worldwide synthesis found that 38% of kelp forests declined over the last five decades [19]. Areas of stability and increase exist, suggesting that in addition to regional climatic trends, local stressors can dominate kelp dynamics. Widespread human activities can impact kelp, including development, agriculture, and forestry [7]. In the last decade, kelp declines and shifts from systems dominated by kelp to turf-forming algae have been documented in Asia, Australia, Europe, North America and South America (reviewed in [20]). Researchers have identified warming [13, 21], eutrophication [22], acidification [23], changes to community structure [18] and sedimentation [24] as contributing factors that often interact. Other known threats include harvest, pathogens, and non-native algal species [25]. As awareness of losses grows along with predictions for future losses associated with climate warming, scientists and managers have identified a pressing need for better baseline information to provide context for past and predicted environmental changes [26, 27] and to determine the extent to which changes are related to human activities.

The Salish Sea is an extensive fjord estuary complex in the northeast Pacific Ocean, spanning the United States and Canada (Fig 1), and two ecoregions [28]. The marine shorelines in the Salish Sea and adjacent exposed coastline support 22 species of kelp [29] with the greatest abundance and diversity found along the western Strait of Juan de Fuca and exposed Pacific Ocean coast [5, 29]. While kelp is less abundant in the inner basins of the fjord complex, the understory kelp *Saccharina latissima* and the canopy-forming bull kelp *Nereocystis luetkeana* (hereafter *Nereocystis*) are common [30–32] where appropriate habitat conditions exist, such as coarse substrates in the shallow subtidal zone for holdfast attachment.

**Fig 1.**
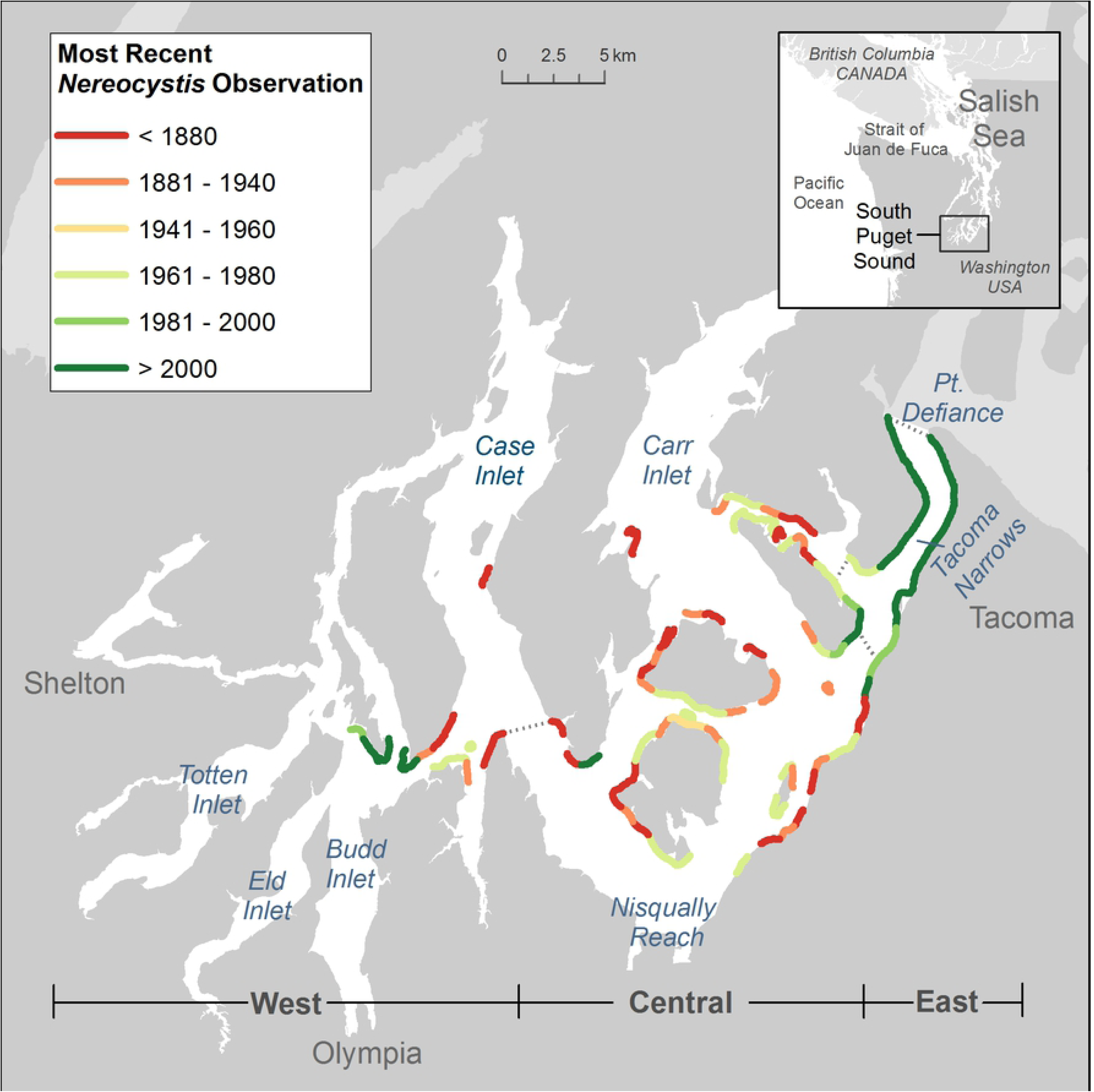
The South Puget Sound study area and three analysis sub-basins (West, Central and East). Marine waters in South Puget Sound (SPS) study area are depicted in white, terrestrial areas are shown in gray and marine areas outside the study boundary are shown in light gray. The -6.1 m bathymetric contour line denotes all shorelines where *Nereocystis* was observed to be present between 1873 and 2018, classified to reflect the year of the most recent observation of presence. Years were binned into 20-year increments, with two bins excluded due to lack of data. The general location of the sub-basins is demarcated at the bottom of the figure, with precise boundaries identified by dotted gray lines on the map.

*Nereocystis* occurs from the Aleutian Islands, AK to Point Conception, CA [33] in a wide range of habitats, from fully wave-exposed to moderately wave-protected shorelines [34]. In the inland waters of the Salish Sea, *Nereocystis* is the sole species of kelp that forms a floating surface canopy. The familiar sporophyte phase of *Nereocystis* is primarily an annual, with a holdfast that attaches to coarse substrates, a long stipe connected to a terminal buoyant bulb, and blades that proliferate on the water surface. The sporophyte has an obligatory alternate phase, a microscopic gametophyte, whose ecology is poorly understood and may be vulnerable to different environmental factors than the macroscopic sporophyte [12, 35, 36]. Limited research has shown that *Nereocystis* populations within Puget Sound are genetically distinct from populations on the exposed coast [37], and they host a distinct microbiome [38].

The Salish Sea was excluded from recent global analyses of kelp trends due to limited scientific studies [6, 19]. In adjacent regions along the open Pacific Ocean coast of Washington and British Columbia, kelp abundance has been highly variable and yet stable over a century-scale [5, 19, 39]. An exception to this general pattern is seen in the eastern extreme of the Strait of Juan de Fuca, where declines occurred along the shorelines furthest within the Salish Sea and most distant from the open coast [5]. Divergent trends in kelp abundance have been noted in adjacent areas associated with distinct local oceanography and human uses in many regions, including British Columbia [40], Nova Scotia [13], Maine [18] and western Norway [22].

Environmental conditions within the Salish Sea differ from areas close to the open ocean in ways that impact kelp performance. There is a general gradient from the ocean into the Salish Sea of decreasing wave exposure, increasing summer water temperatures, decreasing summer nutrient concentrations, and increasing human development [40–43]. South Puget Sound (SPS) is the most distant basin in Puget Sound from oceanic influence and it naturally experiences lower flushing rates and longer water residence times, which make it particularly sensitive to water quality degradation [43, 44]. Human activities impact water conditions in SPS through point and non-point pollution sources associated with regional development in the SPS watershed, as well as in nearby Central Puget Sound [45].

Diverse data sources have noted the occurrence of *Nereocystis* in SPS since European exploration began in the mid-1800s. Historically, kelp surface canopies were charted as an aid to navigation [46]. More recently, surface canopies have been surveyed for environmental monitoring and resource management [47]. In addition to these canopy-focused studies, dive-based ecological studies (ie., [32]) quantified density and other metrics for *Nereocystis* over more limited spatial scales.

Here, we synthesize diverse historical and modern data sources in order to understand spatiotemporal patterns in *Nereocystis* in the SPS. Specifically, we address: 1) What was baseline *Nereocystis* distribution prior to widespread European settlement? 2) During the past century and a half of European development, what were the patterns of *Nereocystis* persistence, loss, and gain? 3) How are historical and recent *Nereocystis* distribution related to environmental characteristics that influence kelp performance, such as water temperature, nutrient concentrations and wave/current energy?

Like other syntheses of diverse historical datasets (i.e., [22, 48]), the primary purpose of our assessment is to describe changes over time. While we cannot draw conclusions about the causes of observed changes, we can place results in the context of regional data characterizing known kelp stressors to draw inferences about some likely stressors. Improved understanding of the historical extent of kelp and patterns of change could support further research into stressors and target restoration and conservation actions. It could also increase our understanding of dynamics in the organisms that rely on these habitats.

## Methods

### Study System

The study area is South Puget Sound (SPS), a 425 km^2^ water body located at the southern terminus of Puget Sound (Fig1), which is part of the larger Salish Sea fjord estuary complex [41, 42, 49]. SPS connects to the northeast Pacific Ocean through a network of basins and sills. While individual basins exhibit distinct oceanographic properties [50], all generally exhibit steadily increasing water residency, stratification, and primary production with distance into the fjord. Tidal currents and estuarine circulation primarily drive water flow and mixing. In SPS, the most intense currents and regular (daily) mixing occur at the Tacoma Narrows, a narrow 1.5 km channel with a shallow sill (45 m) that connects SPS to the rest of Puget Sound. SPS is relatively protected from wave exposure, ranging from semi-protected to very protected in the regional shoreline classification dataset called the ShoreZone Inventory [51].

SPS has complex shorelines composed of islands, passages, and shallow inlets. Due to the area’s glacial origin, gravel, sand, and mixed fine substrates from eroded glacial till and outwash predominate in the intertidal and shallow subtidal zones [30, 52]. Mixed coarse substrates are found along shorelines with strong currents and relatively long fetch. Tideflats of mud and/or sand predominate at the heads of the inlets and other shallow embayments [30, 41].

Archaeological remains show that native people have inhabited the region for more than 12,000 years[53]. European settlement in Puget Sound began in SPS in the 1820s, with establishment of a Hudson’s Bay Company post at Fort Nisqually for fur-trading and agriculture. Population expansion began in the late 1800s, growing from 304 non-native people censused north of the Columbia River in 1849 to 23,955 in 1870 [54]. SPS contained the largest population in the Puget Sound region in 1870, with 1,557 non-native settlers censused in Olympia and Tumwater. After 1870, population growth and density in the Central Puget Sound cities of Seattle and Tacoma outpaced SPS. The Puget Sound region is now extensively urbanized, with a regional population of more than 4 million in 2019 [55]. A number of human activities have impacted natural systems since European settlement began, with lumber production dominating economic activity during the majority of this period. The first sawmill in Puget Sound began operating in 1847 in Tumwater (near Olympia). Initially, trees were felled along the marine shorelines and rivers and floated to mills [56]. In the 1880s, logging activities spread away from waterways to uncut timber accessed by newly built railroad lines. Logging activities expanded rapidly and Washington become the leading state in lumber production in 1905 [57]. Other important economies in SPS have included fishing, agriculture and aquaculture. In recent decades, impacts associated with urbanization predominate. Development has brought extensive nearshore habitat loss and degradation [58], with attendant water quality issues pertaining to anthropogenic nutrient loads and common contaminants from urban, industrial and agricultural runoff.

### Kelp survey data synthesis

We compiled 48 individual data sources that noted the presence or absence of *Nereocystis* in SPS, including peer-reviewed publications, maps, navigation charts, reports and field surveys [30, 32, 59–103]. We limited our analysis to data sources that were recorded in field notes, reports and publications. The datasets spanned from 1873 to 2018 and were produced for a wide range of purposes, including navigation, harvest, resource management, land use planning, and ecological research. The spatial extent of data sources varied from a single location to the entire study area. The format and level of detail also varied widely, including text descriptions of presence or absence at a location, generalized cartographic symbols, detailed delineations of bed perimeter, and phycological studies which examined detailed plant metrics such as density and phenology. When multiple versions of a data source existed, we chose the most detailed field survey in preference to the final product, which was often edited for cartographic presentation. For example, we selected the hydrographic sheets [59] and the accompanying descriptive reports used to create the navigation charts, whenever available. For the Coast Pilot, which was updated gradually over time, records were grouped into three time periods (1889, 1926, 1951) following the methods of Thom and Hallum [67, 70, 86]. In five cases, original data sources were unrecoverable, so we drew data from a compilation of historical data sets [61, 66, 68, 93, 104].

We selected a data synthesis approach to accommodate diverse datasets and also recognize two major known sources of uncertainty that limit the precision of *Nereocystis* canopy observations. First, canopy-forming kelp exhibits high inter-annual variability in extent [5, 105], so a single delineation will generally be a less representative measure of multi-year conditions than for slower growing, longer lived biogenic habitats like coral reefs. Second, tides and currents affect the extent of the *Nereocystis* canopy that is visible on the water surface over short time periods (hours) in this region [106]. These limitations called for a synthesis approach that generalized *Nereocystis* observations to the most comparable format.

We developed a linear model to represent *Nereocystis* presence along the shoreline because the majority of sources depicted the approximate location of *Nereocystis* along the shoreline, rather than precisely delineating the canopy footprint. The linear model captured the common narrow, fringing bed morphology of *Nereocystis* along the SPS shorelines. We selected a -6.1 m (MLLW) bathymetric contour line because it represents a generalized maximum depth of *Nereocystis* beds in SPS and more consistently reflects the linear extent of available *Nereocystis* habitat than intertidal contour lines. A relatively high-quality -6.1 m (MLLW) digital isobath exists, derived from gridded bathymetric data [107].

For all mapped surveys, we transferred information on survey extent and kelp presence from individual data sources to the common bathymetric contour in a geospatial database using ArcMap 10.6.1 [108]. We split the contour line to denote *Nereocystis* presence/absence alongshore, with a minimum mapping length of 3 m for single bulbs (Fig 2). Features were generally one or more orders of magnitude larger than the minimum mapping unit in all datasets except the 2017 synoptic snapshot (S1 Text, S1 Fig). Because many sources constituted general depictions of *Nereocystis* presence, rather than maps, we subsequently further generalized the linear data by summarizing presence/absence over 1-km segments of shoreline. We systematically divided the isobath into 1-km segments, defining 459 segments along the 452 km study area. Of these, 14 segments (3%) deviated from the 1 km length by more than 15%. Twelve segments measured less than 1 km, all occurred along shorelines and islands where the isobath was not an integer multiple of 1 km. Two offshore shoals exceeded 1 km (1.2 and 1.3 km, respectively). For data sources with approximate locations, such as phycological surveys or navigation descriptions from the Coast Pilot, we assigned presence/absence to 1-km segments.

**Fig 2.**
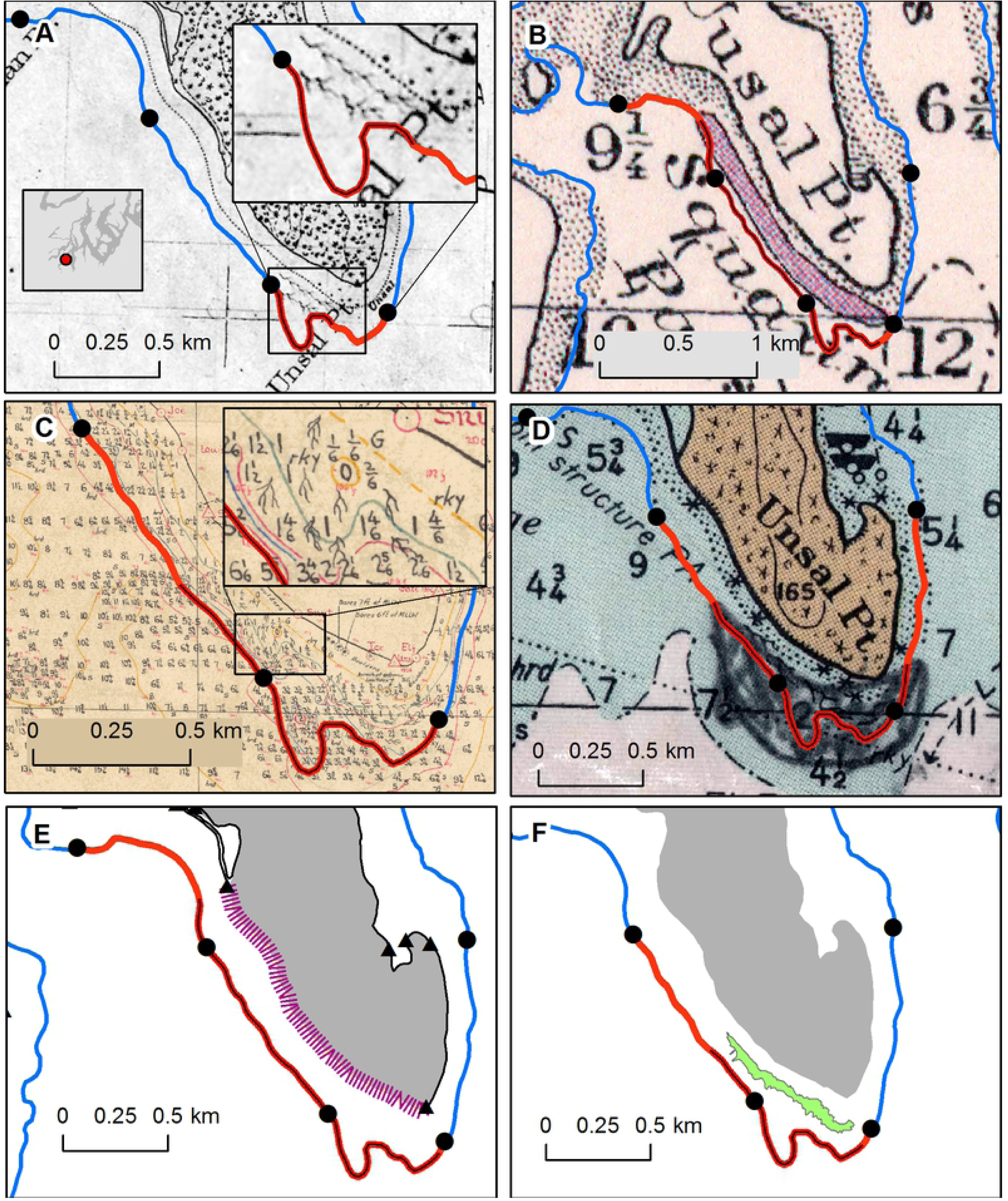
Examples of data sources and compilation methods, showing the -6.1 m (MLLW) isobath contour line overlaid on georeferenced data sources. All maps depict the southern tip of Squaxin Island (sector #1 in Fig 7), scale varies slightly to capture entire area classified with *Nereocystis* for each data source. The isobath (red or blue line) was split into 1 km sections, black circles indicate boundaries. Line colors indicate whether *Nereocystis* was classified as present (red) or absent (blue) in each 1-km segment. The nested black line notes the linear extent of *Nereocystis* delineated in the individual data source. Data sources (see Table 1): (A) topographic sheet #1672 (1878), wavy lines symbolized kelp. (B) kelp harvest map (1911), a polygon symbolized kelp. (C) hydrographic sheet #6198 (1936), wavy lines symbolized kelp. (D) Washington Department of Fisheries field notes (1978), kelp polygon drawn on a chart. (E) The Washington ShoreZone Inventory (1999), floating kelp was noted as patchy or continuous within geomorphologically defined units along the high tide line (black line) with triangles denoting unit ends, purple hatches identify the unit with patchy kelp. (F) DNR habitat map (2017), the perimeter of the floating kelp bed was delineated from kayak at low tide (green polygon) at sites of concern and alongshore extent at other sites.

**Table 1.** Synoptic Nereocystis surveys completed in South Puget Sound.

| Survey Years | Purpose | Reference Year | Data Source Description | Scale |
| --- | --- | --- | --- | --- |
| 1873-1879 | navigation | 1876 | Topographic sheets nos. 1327a, 1327b, 1671, 1672, 1674 1528 [59, 60, 62-65]. Surveyed in the field on plane tables [109]. Geo-referenced maps[110] were aggregated into a synoptic snapshot. (Fig 2A) | 1:10,000 |
| 1911-1912 | harvest | 1911 | Kelp harvest surveys were completed throughout Puget Sound as part of the west coast-wide Fertilizer Investigations [69]. Beds delineated on final maps were wider than actual bed width denoted on preparatory maps. (Fig 2B) | 1:100,000 |
| 1935-1936 | navigation | 1935 | Hydrographic surveys nos. 5931, 6102, 6103, 6104, 6105, 6106, 6107, 6108, 6197, 6198, 6199, 6202, 6203, 6204, 6205 [71-85]. The source data for navigation charts, included field surveys of soundings and aids to navigation. Multiple maps were aggregated into a synoptic snapshot. (Fig 2C) | 1:10,000-1:20,000 |
| 1978 | habitat | 1978 | Washington Department of Wildlife field survey maps[92], annotated in pencil on a paper hydrographic chart, source for the Coastal Zone Atlas[111]. (Fig 2D) | 1:100,000 |
| 1997 - 1999 | habitat | 1998 | WA State ShoreZone Inventory, based on low tide helicopter-based videography [30, 51]. Classified *Nereocystis* alongshore presence as patchy (<50%) or continuous (>50%) within geomorphically-defined linear shoreline units (Fig 2E) | 1:24,000 |
| 2017 | habitat | 2017 | GPS-based small boat survey that noted presence of *Nereocystis* as a line feature along the -6.1 m bathymetry line and as a polygon feature for beds of concern [103]. Single bulbs or aggregations were mapped with a minimum linear width of 3 m. A 10 m threshold between alongshore plants was applied to classify gaps as 'absent'. The 2017 survey was likely to detect more *Nereocystis* than other surveys due to the small minimum mapping unit, the low threshold for *Nereocystis* presence (one bulb), and extensive field effort. (Fig 2F) | 1:12,000 |

We evaluated the two alternative approaches (linear extent vs 1-km segments) for mapped data sources, the results were similar (S1 Text, S2 Fig). We concluded that the 1-km segment scale was superior because it generalized the highly variable levels of resolution among surveys, and it also allowed for inclusion of data sources that were not precisely located. The limitation of 1 km shoreline segment approach was that it could not detect subtle changes in distribution, such as changes in the extent of *Nereocystis* within a segment. However, because differences among survey methodologies generally precluded identification of subtle changes, the benefit of including more data sets outweighed the cost of decreased precision and allowed more temporal variability across the domain to be evaluated.

We recorded 3,232 instances of *Nereocystis* presence/absence between 1873 and 2018 at 1-km segments. At segments where *Nereocystis* occurred at least once, the number of presence/absence observations per segment over the entire time period ranged from 6 to 14, with a median of 8. Fewer data sources existed in the heads of embayments where *Nereocystis* never occurred, leading to a slightly lower study area-wide median of seven observations per segment.

We used a series of metrics to assess spatiotemporal patterns in *Nereocystis* distribution. First, we selected six synoptic datasets, separated by 19 to 44 years, that each comprehensively assessed the study area over a limited time period (Table 1). For simplicity, we refer to each synoptic snapshot with a single year from the mid-point of the surveys. The six synoptic snapshots employed the majority of the presence/absence observations (2,239 of 3,232 observations, or 69%). The earliest synoptic snapshot was completed in the 1870s, which constitutes a baseline of conditions early in the history of European settlement in the Puget Sound region. We assessed changes in kelp distribution throughout SPS from this baseline to recent conditions by comparing the number of 1-km segments with *Nereocystis* present and proportion of segments per sub-basin with *Nereocystis* present in the 1870s and 2017. These two surveys were relatively detailed and based on extensive field surveys (Table 1).

While the synoptic snapshots allowed for comparison of distinct time points, they excluded 31% of the segment-scale observations, including highly detailed phycological studies and dive surveys. We employed the entire pool of observations to further explore patterns in the most recent *Nereocystis* occurrence per segment and overall persistence. We defined persistence as the proportion of all data sources at each 1-km segment that noted *Nereocystis* presence. Because the data showed a shift from presence to absence at many segments after the 1978 synoptic survey, we calculated persistence before and after 1980.

To compare patterns of kelp abundance and distribution over time within sub-areas, we divided the SPS basin into 3 sub-basins (Fig 1). The sub-basins partitioned the area along a gradual seasonal gradient in water temperature and nutrient concentrations [44, 112].

We summarized patterns in presence/absence from all surveys by lumping segments where *Nereocystis* ever occurred into contiguous sectors less than 10 km in length. Sector boundaries considered geomorphological features, such as headlands, and oceanographic features, such as fetch and aspect. We defined 30 sectors, a tractable number of locally recognized geographic landmarks for local management and research. Because initial analyses showed strong declines in *Nereocystis* in many sectors, we summarized presence to be the occurrence of any *Nereocystis* within each sector.

### Water temperature, salinity and nutrient concentration data

We explored the relation of surface water temperature, salinity and nutrient concentrations to observed kelp distribution by characterizing long-term conditions from mid-channel water quality monitoring stations and recent conditions from nearshore stations.

#### Long term, mid-channel conditions

We described long term water properties with publicly available data from four mid-channel water quality stations, sampled monthly by the Washington State Department of Ecology [113]. We restricted our analysis to the top five meters of the water column characterized in continuous vertical profiles measured with a conductivity, temperature and depth (CTD). We summed nitrate, nitrite and ammonium concentrations to represent dissolved inorganic nitrogen (DIN) at the two shallowest reported depths for nutrients (0 m and 10 m). Monthly values for each parameter at each depth for all available years were reduced to a representative annual pattern using locally estimated scatterplot smoothing (LOESS). More than two decades of monthly data exist at three of the stations, DNA001 (1989-2016), NSQ002 (1996-2016) and GOR001 (1996-2016). At NRR001, data is limited to three years (1989-1991).

#### Recent nearshore conditions

We characterized recent nearshore water column properties along an axis from the entrance to SPS at Tacoma Narrows to the most distal documented kelp forests using data collected at seven nearshore sites monthly from September 2017 to August 2018. This axis encompasses a known environmental gradient [113], and sites were placed near historical and recent *Nereocystis* beds (Fig 3). All sites were surveyed on the same day, within two hours of solar noon, during low tide and low current periods. A sampling station was established in the center of each site along the -6.1 m (MLLW) bathymetric contour. A weighted *SonTek Castaway®-*CTD measured temperature and salinity. We calculated mean salinity and temperature for the top 5 m of the water column. From March to September 2018, field-filtered water samples were also collected to measure DIN concentrations at four sites as time permitted (Squaxin Island, Devil’s Head, Day Island, Salmon Beach). From May to September, samples were taken from each site at 0.25 m and 4 m depths in order to assess possible surface water stratification during late-spring and summer. In March and April, only 1 sample was collected at 4 m depth. An acid washed 60 mL syringe with an attached 0.45 µm cellulose acetate filter was filled with water directly from a Van Dorn sampler. A small amount of water was filtered through the syringe to rinse the syringe and syringe filter before rinsing an acid washed 60 mL high density polyethylene bottle with filtrate. The bottle was then filled with filtrate before being placed immediately in a cooler on ice and transported to the Evergreen State College laboratory where they were frozen (−10° C) for later transport to the University of Washington’s Marine Chemistry Lab for total dissolved nutrient analysis using spectrophotometric methods.

**Fig 3.**
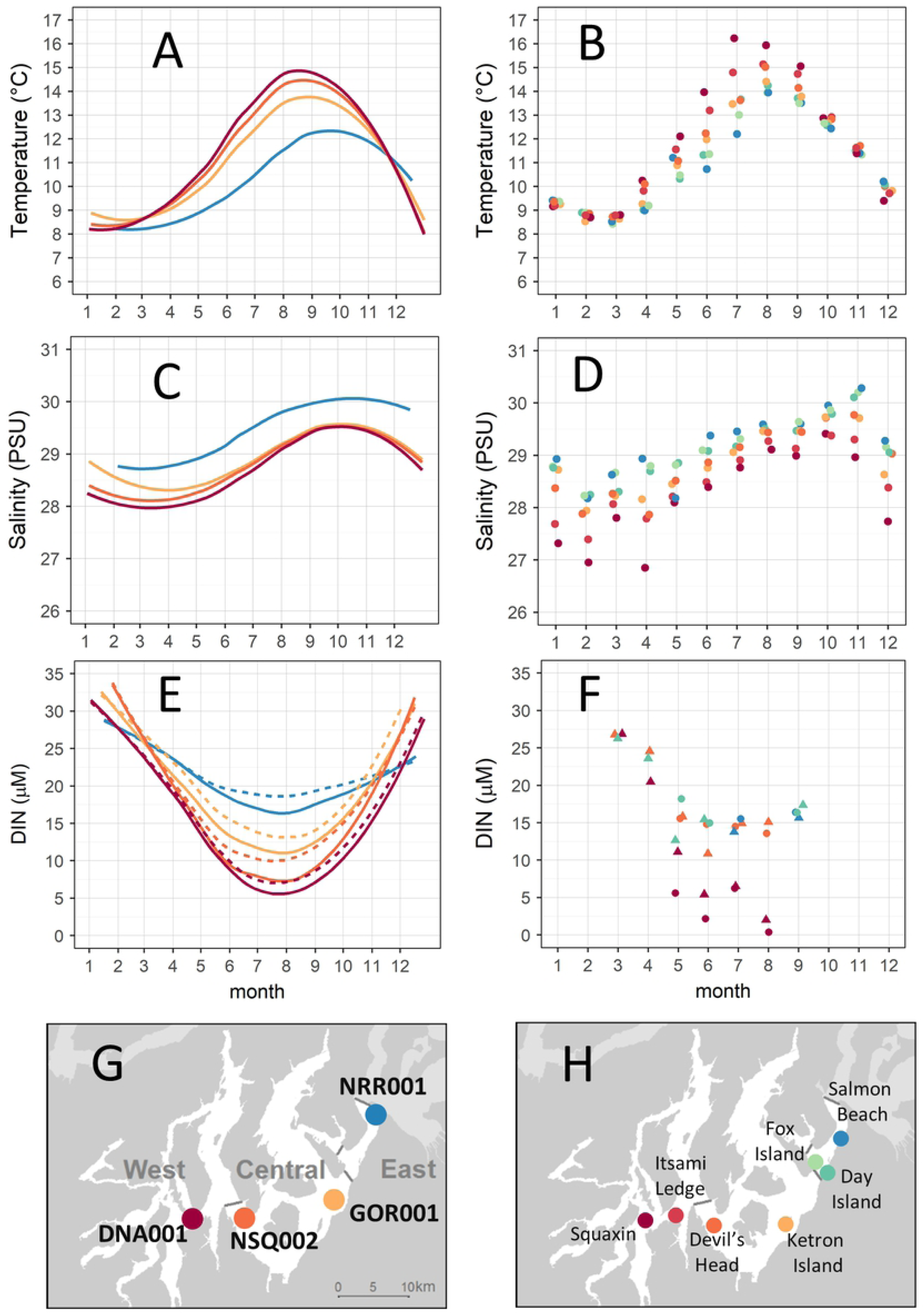
Monthly water characteristics at mid-channel long-term monitoring stations (left) and nearshore stations (right). (A, B) mean water temperature to 5 m depth. (C, D) mean salinity to 5 m depth. (E) DIN concentration at mid-channel stations at depths of 0 m (solid line) and 10 m (dashed line). (F) DIN concentration at nearshore stations at depths of 0.25 m (point) and 4 m (triangle), with data slightly offset horizontally for visibility. Site locations for (G) mid-channel stations and (H) nearshore stations. Mid-channel long-term monitoring station data (left) represent a cubic spline curve fit to mean values from more than 2 decades of sampling for all stations except NRR001 (3 years). Nearshore stations (right) were sampled monthly at -6.1 m (MLLW) between September 2017 and August 2018.

### Wave and current exposure characterization

We used modelled current velocity and wave heights to explore the relationship between these physical factors and spatiotemporal patterns in *Nereocystis*. We limited these comparisons to segments were *Nereocystis* was observed at least once (n=120).

We characterized current speed using modelled surface water velocity data from a Salish Sea circulation model [43, 114, 115]. We used data from the surface layer, which represents the top 3% of the water column, from model year 2014. We calculated the annual average of the maximum daily flow velocity (m/s) based on the flow velocities in the x and y directions at each model node. We then summarized velocity with a single value for each 1 km shoreline segment by selecting the value from the closest model grid point. Median distance between model node points and the corresponding shoreline segment was 91 m.

We characterized wave energy using average annual maximum wave height data developed by the Washington Coastal Resilience Project using the numerical wave model SWAN (Simulating WAves Near Shore) [116]. The model generated a hindcast of hourly wave conditions across the Salish Sea over a 60-year period between 1950 and 2010. Modeled values were sampled along the -10 m (NAVD88) bathymetric isobath for SPS and quantified the average annual maximum wave height over the 60-year hindcast. The hindcast utilized the 12-km Weather Research and Forecasting historic reanalysis of the Pacific Northwest [117, 118] which was found to represent the spatial patterns of extreme wind events well, but bias wind speeds slightly lower (∼1 m/sec) than observed winds over water. Because of this slight bias and because significant wave heights characterize the upper 33% of the wave-height distribution, the wave heights reported here are likely underestimates. We summarized the wave data every 1-km alongshore by selecting the wave model grid point closest to each 1-km segment. Median distance between the model point data and the corresponding shoreline segment was 3.4 m.

### Statistical analyses

All data analyses were performed in ArcGIS version 10.6.1 [108] and R 3.6.0 [119]. Data analysis of kelp extent focused on describing spatial and temporal patterns over time because methods varied widely among data sources and replicate samples did not exist.

We tested if temperature was different among sites during summer (June to September) or winter (November to February) using two mixed effects models (one for each season) with a random factor of month (temperature ∼ site, random = 1|month), with the R packages “nlme”[120] and “emmeans” [120]. We tested if residuals were normally distributed using qqplots and Shapiro-Wilk tests and visually assessed model output for patterns in normalized residuals.

We tested for differences among sub-basins in current speed and wave height at sites where *Nereocystis* was observed at least once using Welch’s ANOVA to accommodate unequal variance and unequal sample sizes, with the R package “userfriendlyscience” [121]. The wave data were normally distributed, and the current data were log transformed to achieve a normal distribution.

## Results

### *Nereocystis* extent declined and spatial distribution shifted

Based on all available data sources, *Nereocystis* occurred at least once along 26% of the SPS shoreline (120 of 452 1 km shoreline segments) between 1873 and 2018. *Nereocystis* never occurred in the extreme reaches of any inlets (Fig 1). When aggregated, the six synoptic snapshots recorded a similar total *Nereocystis* extent over the entire time period as the complete collection of data sources (115 segments and 120 segments, respectively). The extent of *Nereocystis* classified in synoptic snapshots fell into two distinct groups (Fig 4): more than 60 segments were identified with *Nereocystis* in 1876, 1935 and 1978 (65, 68, and 65 segments, respectively). In contrast, *Nereocystis* occurred in one-third as many segments in 1911, 1998, and 2017 (16, 26, and 25 segments, respectively). Differences among datasets in total extent likely reflect both changes in kelp distribution and methodological differences among surveys. The 1911 estimate was the lowest, and it starkly contrasts the high estimates before (1876) and after (1935). The 1911 estimate could have marked a minimum in *Nereocystis* extent, however it is also likely that the project objective to identify beds with harvest potential for fertilizer resources led to identification of a limited number of large, accessible beds rather than an exhaustive survey (Table 1, S1 Text, S1 Fig, Results section). The most recent estimates from 1998 and 2017 suggest that *Nereocystis* extent was dramatically restricted in recent years to one-third of the extent in 1876, 1935 and 1978.

**Fig 4.**
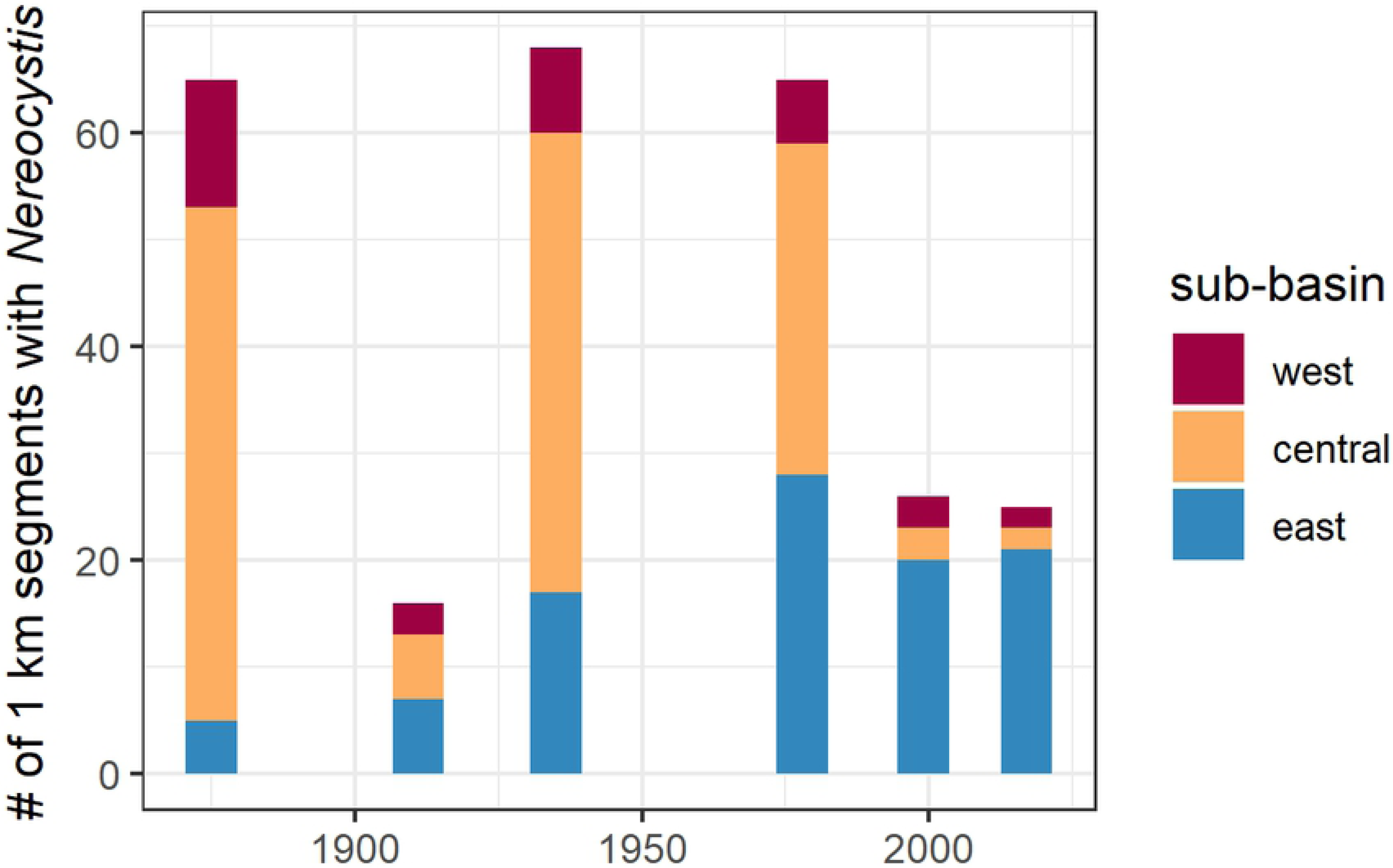
Number of 1-km segments with *Nereocystis* present between 1876 and 2017, based on six comprehensive snapshot surveys, summarized over three sub-basins. Recent estimates (1998 and 2017) are dramatically reduced relative to estimates in 1876, 1935 and 1978. The 1911 estimate could represent a low point in kelp extent, but likely reflects methodological differences in survey methods (harvest study).

The six synoptic snapshots showed a marked shift in the spatial distribution of kelp forests among sub-basins over time (Fig 4). The Central sub-basin had the most shoreline with *Nereocystis* in 1876, 1935, and 1978 (74%, 63% and 48%, respectively) and a third of the total in 1911 (38%). In contrast, the Central basin only contained a tenth of the total extent in the two most recent surveys (12% in 1998 and 8% in 2017). The West sub-basin generally contained a smaller proportion of the total shoreline with *Nereocystis* than the Central sub-basin, 8-19% over the entire time period. Proportional decreases in Central and West corresponded to increases in the East, where the proportion increased from 8-44% during the earliest four surveys to 77-84% in the two most recent surveys.

The 1876 synoptic survey constitutes the oldest known comprehensive temporal baseline, surveyed early in the history of European settlement in the region and SPS. When compared to the most recent synoptic survey in 2017, *Nereocystis* extent decreased by 62% throughout the SPS study area, from 65 to 25 1-km segments. The most extreme losses occurred in the Central sub-basin, where kelp decreased by 96%, followed by the West sub-basin, where *Nereocystis* decreased by 83%. In the Central sub-basin in 2017, *Nereocystis* occurred in only two isolated segments, while in the West sub-basin *Nereocystis* was confined to a single bed that spanned two contiguous segments. In contrast, the linear extent in East sub-basin more than tripled to 21 segments.

### Most recent occurrence of *Nereocystis* and persistence over time

Analysis of all data sources provided similar results as the synoptic snapshot surveys, and greater detail on patterns of persistence at segments and the timing of the most recent occurrence of *Nereocystis*. The East sub-basin contained the greatest proportion of recent occurrences; *Nereocystis* occurred at 72% of the segments since 2000 and at all segments since 1960 (Fig 1 and Fig 5). In contrast, in the Central and West sub-basins, at the majority of segments the most recent *Nereocystis* occurrence was prior to 1980 (89% and 63%, respectively).

**Figure 5.**
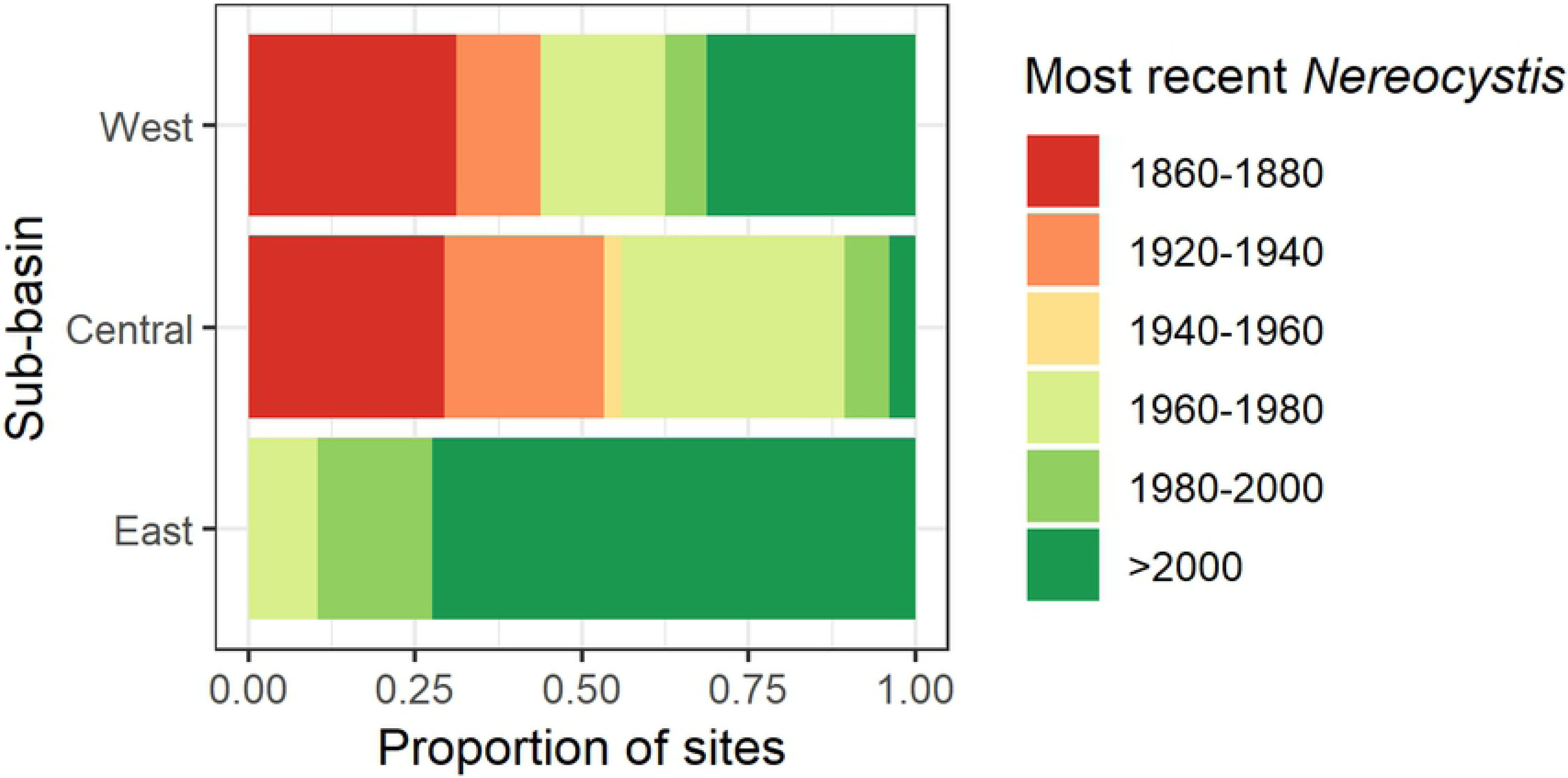
Year of most recent *Nereocystis* occurrence within each sub-basin. Bar plots show the proportion of segments within each sub-basin. Years were binned into 20 year increments, with two bins excluded due to lack of data. All 1-km segments where *Nereocystis* occurred at least once during the entire study period were included (n=120).

Persistence, measured as the proportion of data sources that noted *Nereocystis* presence within each 1-km segment, ranged from 0.1 to 1, with a median of 0.3. There was a marked difference among sub-basins in *Nereocystis* persistence before and after 1980 (Fig 6). Before 1980, persistence was similar in West and Central (median values of 0.27 and 0.22, respectively), and slightly higher in East with a median of 0.55. After 1980, median persistence dropped to 0 in the West (13 of 16 segments decreased) and 0 in the Central sub-basins (73 of 74 segments decreased) and remained stable in the East sub-basin (median value of 0.60, 9 of 28 segments decreased). After 1980, most segments with high persistence were restricted to the East sub-basin (Fig 7). In the Central sub-basin, two spatially isolated beds remained, along with beds along the approach to the Tacoma Narrows (Fig. 7). In the West sub-basin, the bed located off the southern tip of Squaxin Island remained highly persistent throughout both time periods (sector #1 in Fig 7).

**Figure 6.**
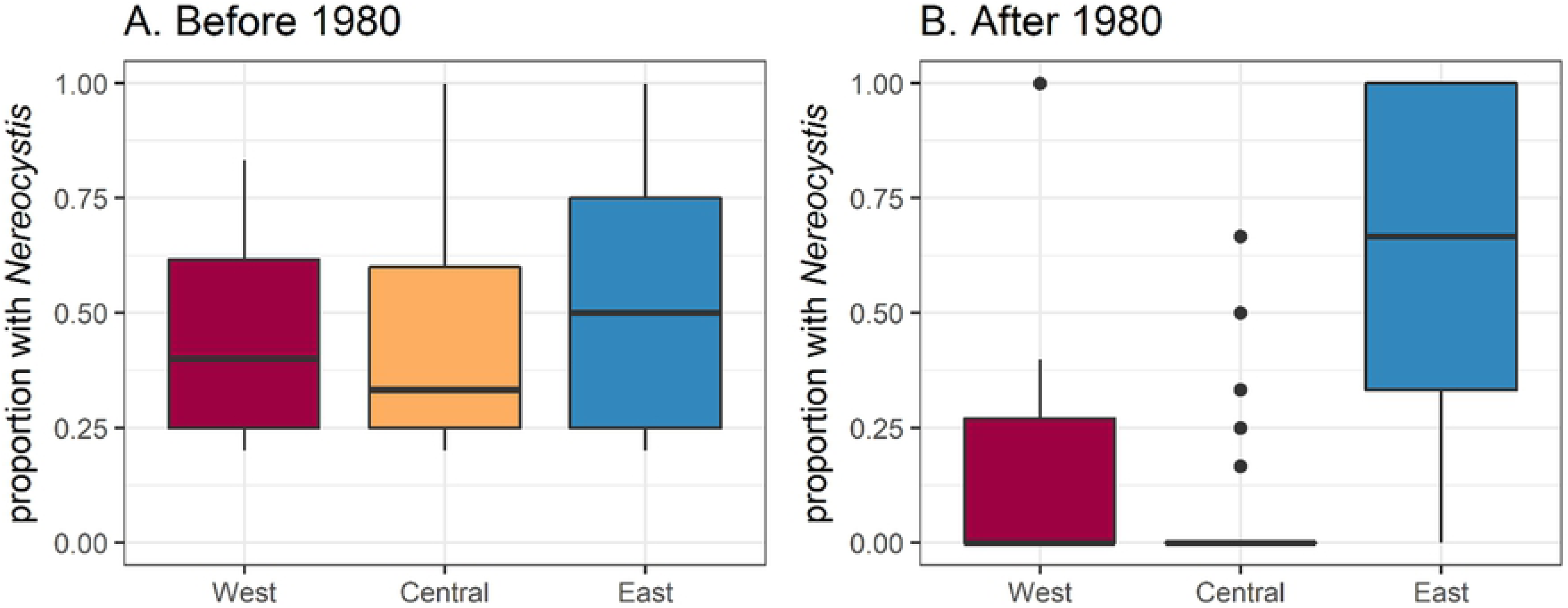
Distribution of *Nereocystis* persistence at 1-km segments (A) before 1980 and (B) after 1980. Persistence was calculated as the proportion of all observations in each segment with *Nereocystis* present within each time period. All 1-km segments where *Nereocystis* occurred at least once in either time period were included (n=120).

**Figure 7.**
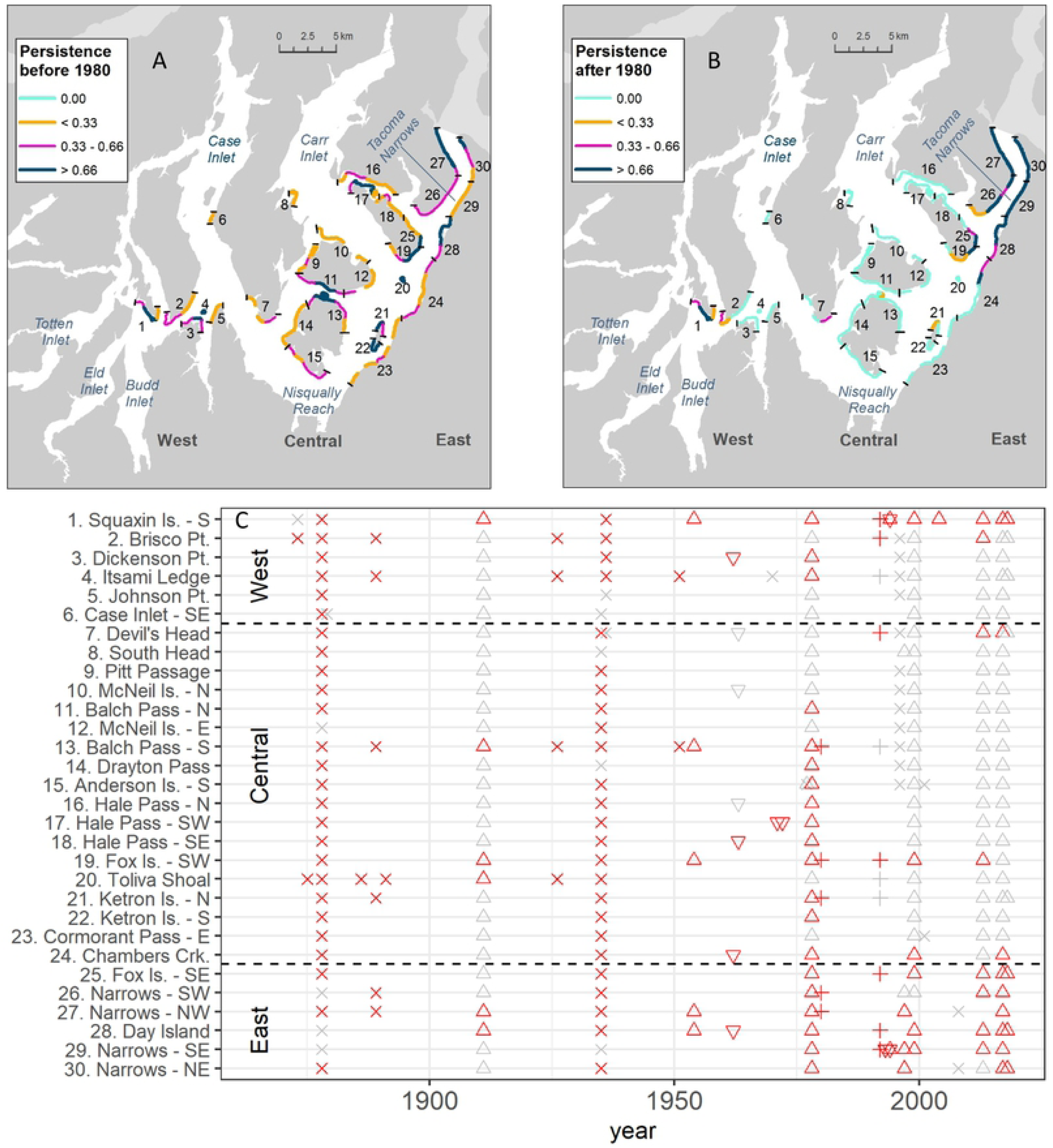
Persistence of *Nereocystis* at segments and sectors (groups of segments). Maps show persistence at segments (A) before 1980 and (B) after 1980. Persistence was calculated as the proportion of all observations with *Nereocystis* present within each time period. Segments where *Nereocystis* never occurred are not shown). The maps also show groups of adjacent segments aggregated into shoreline sectors (<10 km in length), with numeric identifiers for each sector and black tick marks delineating boundaries. (C) identifies all sectors by geographic name and number and summarizes all observations as presence/absence for each sector by survey year. Color denotes presence (red) or absence (grey). Shapes represent dataset type (‘x’ for navigation chart, triangle for kelp/habitat survey, inverted triangle for phycological survey, + for fish survey).

Generally, sectors in the West and Central sub-basins most often exhibited a pattern of presence in early years and absence in recent decades, with a common transition following the 1978 synoptic survey (Fig 7C). One widespread exception to this temporal pattern occurred in 1911, when little *Nereocystis* was recorded overall. This exception could be an artifact of project intent to identify economically viable beds for harvest (Table 1), or it could reflect response to intensive logging along the shoreline during early settlement or it could reflect a natural minimum in *Nereocystis* distribution.

Detailed site-scale studies confirmed the presence of *Nereocystis* at many segments prior to the shift observed in large area mapping projects around 1978. Phillips [88] documented a fringing *Nereocystis* bed during a dive survey in 1962 at Dickenson Point (sector #3 in Fig 7) [88]. One of the authors (Mumford) remembers *Nereocystis* beds at Dickenson Point and nearby Itsami Ledge (sectors #3 and #4 in Fig 7) in the 1970s, with the last occurrence of scattered plants at both sectors in the early 1980s. Hodgson and Waaland [91] recorded *Nereocystis* in a multi-year phycological study in 1971 and 1972 at Hale Pass SW (sector #17 in Fig 7). In the Tacoma Narrows where *Nereocystis* has been noted to be present in recent years, Maxell and Miller [32] confirmed presence in 1993 and 1994 in a dive-based phycological study (sector #28 in Fig 7). More recently, a second distinct shift occurred in 2017 and 2018 [103], when *Nereocystis* disappeared from Brisco Pt, Devil’s Head and Fox Island SW (sectors #2, #7 and #19, respectively). These recent losses further restrict the recent distribution of *Nereocystis* in SPS (Fig 8). None of the beds that disappeared have returned. While *Nereocystis* beds persist at Squaxin Island and Fox Island SE (sector #1 and #25, respectively), intensive monitoring documented declines in bed area between 2013 and 2018 [103].

**Figure 8.**
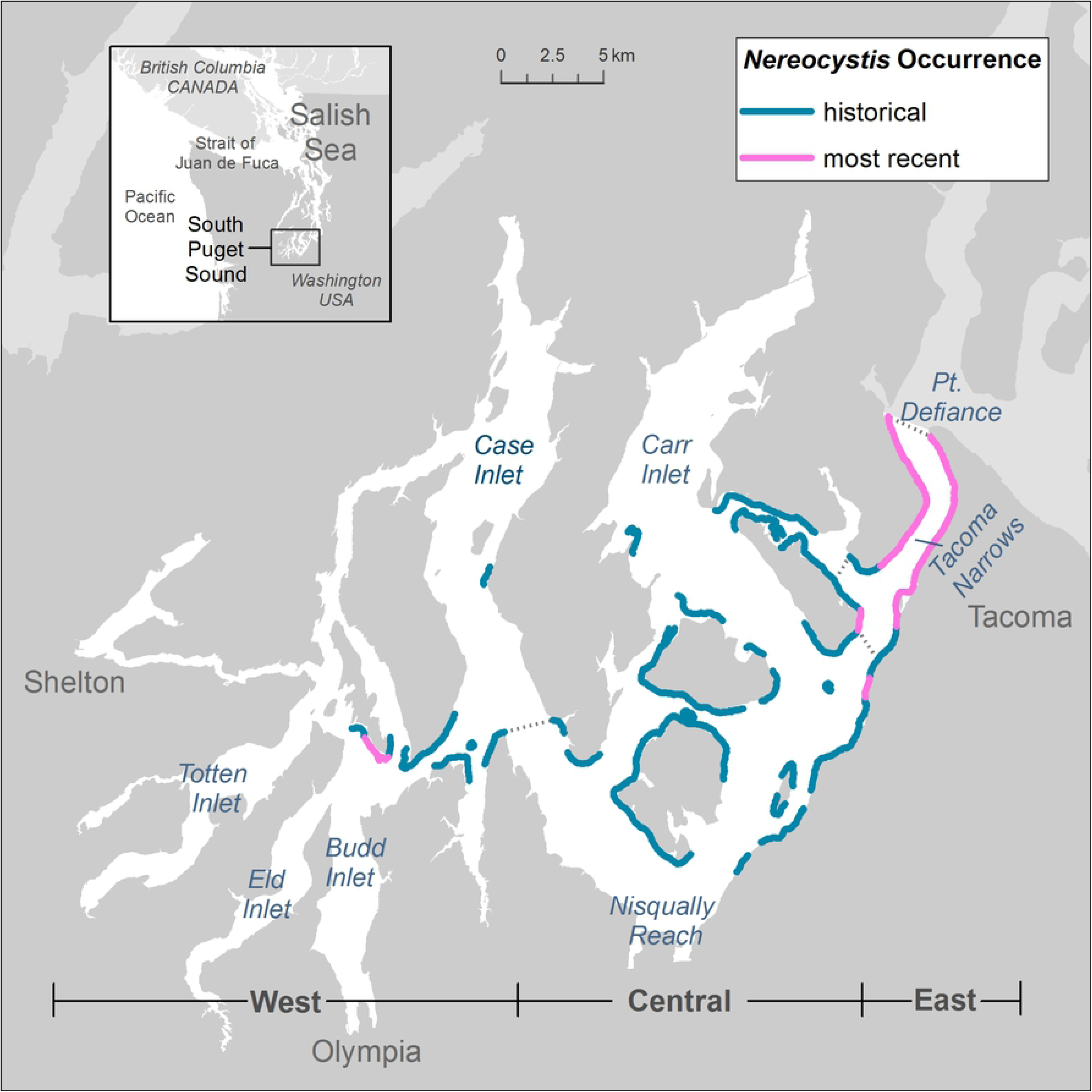
Most recent estimate of *Nereocystis* distribution in SPS. The -6.1 m bathymetric contour line denotes all 1-km segments where *Nereocystis* was observed to be present between 1873 and 2018. Pink denotes segments where *Nereocystis* was present during most recent observation (2017 or 2018) while blue denotes segments where *Nereocystis* was present in the past and absent in the most recent observation.

### Sea surface temperature and nutrient data show strong gradient

Water temperature, salinity and nutrient concentration data displayed strong seasonal and spatial patterns that were similar at the long-term mid-channel stations and the nearshore stations sampled in 2017-2018 (Fig 3). For the winter nearshore temperature data, the likelihood ratio test indicated that including site did not improve the final mixed effects model (L = 4.7, df = 1, p = 0.034), indicating that average winter water temperature did not differ among sites. In contrast, nearshore summer water temperatures varied markedly among sites (L = 52.52, df = 1, p < 0.001). Post-hoc tests confirmed that the magnitude of differences in summer temperature increased with geographic distance between sites. Minimum annual temperatures (8°C) occurred during February/March, with less than 1°C difference among all stations. From March to October, the warmest water consistently occurred in the West sub-basin at adjacent locations in Dana Passage (DNA001 mid-channel station) and Squaxin Island (nearshore station), with slightly higher measurements at nearshore stations. The highest overall water temperature recorded at a nearshore station occurred at Squaxin Island in July 2018 (16.2 °C). The coolest spring-fall temperatures consistently occurred in the East sub-basin at adjacent locations in the Tacoma Narrows (NRR001 mid-channel station) and Salmon Beach (nearshore station). At Salmon Beach, nearshore temperature peaked at 13.9 °C in August 2018. Central sub-basin water temperatures fell midway between the extremes measured at Tacoma Narrows and Dana Passage at all of the mid-channel and nearshore stations, with a consistent gradient of values increasing with distance from the Tacoma Narrows. The water column at all nearshore stations was well mixed even in July during peak annual temperatures, with less than 0.5 °C range per cast between the surface and 5 m depth (S3 Fig).

Salinity ranged from 28 – 30 PSU (long-term curve-fit) at mid-channel stations, with similar values and annual patterns at nearshore stations. Salinity was higher in the summer and late fall, a common pattern in the region associated with annual rainfall and freshwater input cycles. Extreme salinity values occurred at the geographic extremes of Tacoma Narrows and Dana Passage. Nearshore salinity ranged from 27.1 PSU in February at Squaxin Island to 30.3 PSU at Salmon Beach in November.

DIN concentrations at mid-channel stations were high in the winter months at all stations; generally, 25 µM and greater at depths of 0 m and 10 m (Fig 3). Values diverged throughout the spring, with pronounced differences among sites from May to October. Concentrations were slightly higher at depth. A strong spatial gradient emerged, with decreasing concentrations into SPS and the most extreme drawdown of nutrients at Dana Passage, where the long-term mean fell below 10 µM at both 0 m and 10 m depth from June to September. Nearshore DIN concentrations showed a similar pattern for all months with data. Nearshore concentrations were indistinguishable at stations sampled during March 2018, ranging from 26 - 26 µM at 4 m depth. Concentrations at Squaxin dropped every successive month until August, reaching the lowest concentration measured at any site in August 2018 (0.4 µM at 0.25 m depth). In contrast, nearshore DIN concentrations at the other stations never dropped below 10 µM.

### Recent *Nereocystis* observations predominated in high current areas

Average maximum daily current velocities ranged from 0.14 to 2.59 m/s (median 0.52 m/s) at segments where *Nereocystis* was observed at least once during the entire study time period (Fig 9). Current velocities differed among sub-basins (Welchs ANOVA test: F(2, 116) = 61.85, p < 0.001). The East sub-basin experienced significantly larger current velocities than the West and Central sub-basins (Games Howell post hoc test, p<0.001), where median velocity (1.63 m/s) was approximately 4 times larger than in the West (0.42 m/s) and Central (0.48 m/s) sub-basins. Comparison of historical *Nereocystis* observations with current velocity data showed all of the segments where *Nereocystis* has not been observed since 1980 or earlier experienced current velocities of 1 m/s or less. After 2000, the majority of segments with *Nereocystis* were restricted to shorelines with mean maximum daily currents above 1 m/s, which were predominantly located in the East sub-basin. However, notable exceptions exist to this pattern; the remaining segments with *Nereocystis* in West and Central sub-basins range in current velocity from 0.31-0.82 m/s.

**Figure 9.**
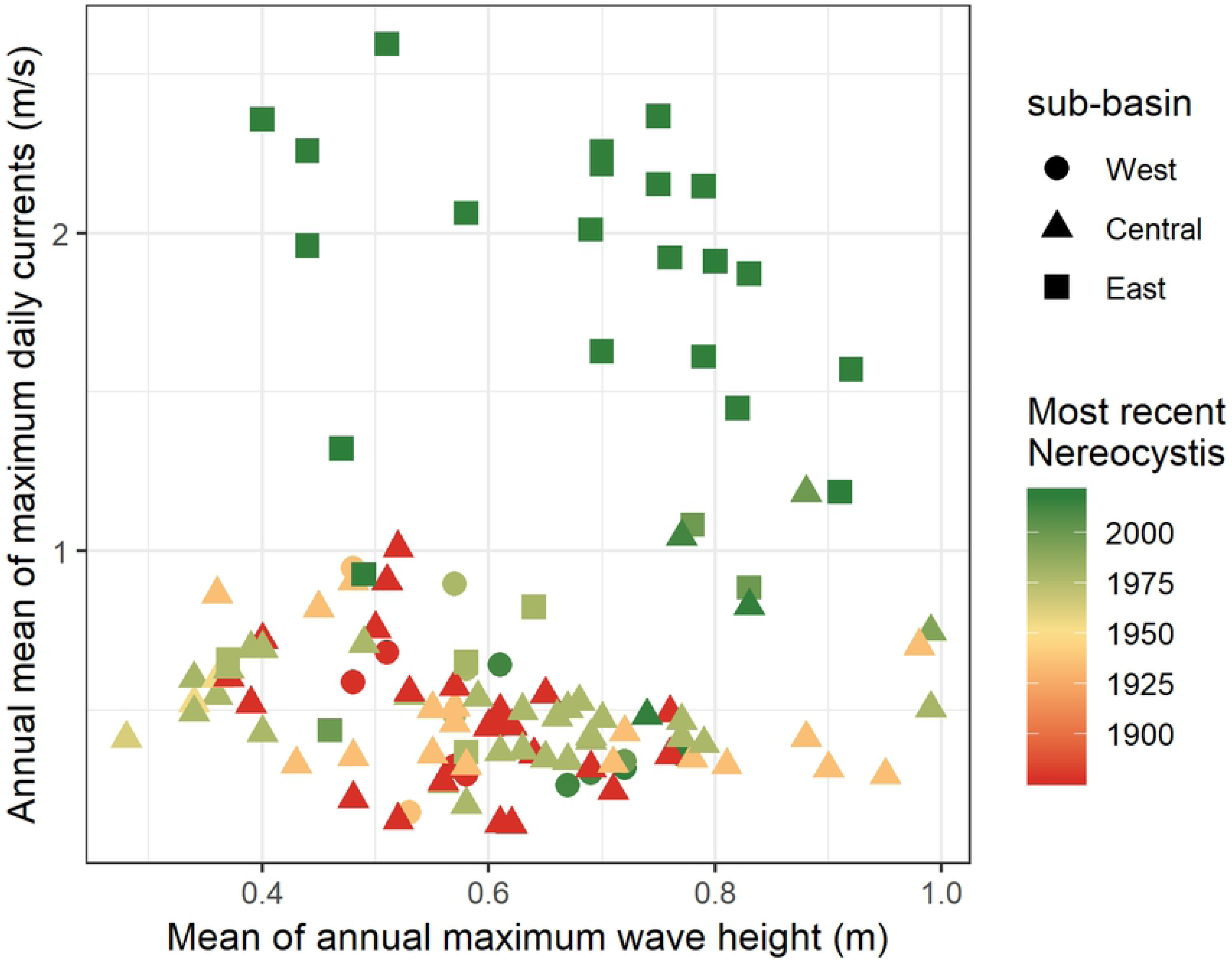
Current and wave exposure at 1-km segments with *Nereocystis*. The annual mean of maximum daily current speed (y-axis) is derived from a 2014 model run of the Salish Sea Model [110, 111]. Values represent the average maximum daily current velocities in the upper 3% of the water column at the nearest model node. Average annual maximum wave height along 1 km shoreline sections was estimated with model data from 1950 to 2010 developed by the Washington Coastal Resilience Project [112]. Segments are coded by sub-basin and the most recent year that *Nereocystis* was observed.

Average annual maximum wave height ranged from 0.28 m to 0.99 m (median 0.62 m) at segments where kelp was observed at least once during the entire study time period throughout SPS. Wave height did not differ among sub-basins (Welchs ANOVA test: F(2, 116) = 0.7, p =0.50) but displayed high spatial variability dependent on shoreline orientation. Locations facing South/North recorded higher annual maximum wave heights, indicative of the dominant wind directions in Puget Sound. Median values within the sub-basins ranged from 0.58 – 0.70 m. No patterns were evident over time in the range of wave heights at segments where *Nereocystis* was observed (Figure 9).

## Discussion

### Baseline shows major *Nereocystis* losses and shift in distribution

This study established a historic baseline for *Nereocystis* distribution in SPS during the 1870s, early in the period of European settlement in the region. We described major losses from that baseline in both extent and distribution over 145 years. The most extreme decreases occurred in the Central and West sub-basins; the most recent dataset identified only one isolated location with *Nereocystis* remaining in each of these sub-basins. In contrast, the East sub-basin appeared stable or increasing. Many of the observed losses in the West and Central sub-basins have persisted for four decades or longer, which demonstrates they are not associated with inter-annual variation.

The observed trend of *Nereocystis* decrease in SPS over 145 years contrasts sharply with findings along the Strait of Juan de Fuca, at the entrance to the Salish Sea. There, kelp forest area generally remained stable over the last century, except along the eastern boundary—the area farthest from oceanic influence and closest to anthropogenic development [5]. This contrasting pattern of adjacent sub-regions experiencing loss and stability has occurred in other locations globally [13, 18, 22, 40].

### Conditions associated with observed patterns in *Nereocystis*

While data on long-term trends in temperature and nutrient concentrations were not available, data from recent decades characterized general conditions. Elevated temperatures and low nutrient concentrations occurred in tandem in SPS, a pattern seen in upwelling systems along the exposed coast and in other areas within the Salish Sea [14, 122–126]. In SPS, summer temperatures were considerably higher and nutrient concentrations were considerably lower compared to the exposed coast and Strait of Juan de Fuca where upwelling and ocean mixing drive water column properties [127]. Many studies have shown that kelp is sensitive to high temperatures and low nitrogen. An 18-year study concluded that temperature increases from a thermal outfall were associated with the virtual disappearance of *Nereocystis* [128]. In British Columbia, elevated temperatures have been associated with lower abundance of *Nereocystis* [105] and other kelp species [40]. In SPS, *Nereocystis* appeared stable in the East sub-basin, where temperature did not exceed 14.0 °C, a threshold used in southern California for compromised kelp performance [129]. In contrast, summer temperatures at nearshore stations in the West and Central sub-basins consistently exceeded this threshold, and additional sampling within the Squaxin Island kelp forest documented even higher temperatures than at the nearshore stations, ranging from 17 °C to 20 °C [103]. These temperature maxima approached or exceeded thresholds for decreased resilience of sporophytes and increased zoospore mortality [36, 130, 131].

Nitrogen requirements for *Nereocystis* are not defined, yet data suggest that low summer concentrations in recent decades may be impacting *Nereocystis* performance in portions of SPS. The authors observed that the *Nereocystis* blades at Squaxin Island were thin, relatively short and shredded. In field studies, concentrations of 10 µM were associated with thicker blade tissues and a lower rate of blade erosion in *Macrocystis* sporophytes [132]. Laboratory studies showed increased performance in microscopic stages of *Nereocystis* associated with DIN increases from 1 to 15 µM [36]. Nitrogen requirements may be greater during periods of rapid growth or elevated temperatures [133]. In Puget Sound, anthropogenic inputs are a major local source of DIN [134]. However, worldwide research predicts that elevated anthropogenic nutrient loads damage kelp performance by stimulating growth of phytoplankton [43] and nuisance algae, and introducing particulates and other pollutants and contaminants [20, 135]. While long-term trends are not well understood, anthropogenic DIN inputs in Puget Sound have altered dissolved oxygen levels and algal biomass [43], and the nutrient balance appears to have shifted in recent decades with potential impacts to species composition and material cycling [136, 137].

Notable exceptions exist to the general pattern of *Nereocystis* losses in sub-basins with higher temperatures and lower nutrients. The innermost bed at Squaxin Island remained highly persistent despite the poorest measured environmental conditions. This exception demonstrates that *Nereocystis* can persist in elevated temperature and low nutrient conditions. The authors believe that the long fetch from prevailing, year-round southern winds and extensive appropriate shallow subtidal habitat contribute to *Nereocystis* persistence. However, even there, more intensive studies found that total bed area and maximum depth decreased between 2013 and 2018 [103]. Another important exception to the general pattern was the widespread *Nereocystis* loss in the Central sub-basin, where temperature and nutrient concentrations were intermediate. These exceptions demonstrate that additional factors outside the scope of this study contributed to trajectories of kelp persistence or loss.

Many studies have demonstrated that hydrodynamic exposure to waves and currents influences kelp dynamics directly and indirectly. Along exposed coastlines, physical disturbance through extreme wave events can drive kelp mortality [15]. In areas of low water motion, the boundary layer can limit the capacity of kelp to acquire nutrients and eliminate waste products [7, 138]. In relatively sheltered environments in other regions, wave exposure metrics were positively correlated with greater kelp performance and negatively correlated with elevated temperature [40]. SPS has a relatively protected wave environment, with short-period waves (<5 sec) that have significantly less energy than long-period ocean swell in other habitats where *Nereocystis* occurs. In SPS, current velocity is the primary source of daily water turbulence, and strong tidal currents lead to lower temperatures and higher nitrogen levels through mixing, especially at the Tacoma Narrows [49]. These factors could explain the observed pattern of *Nereocystis* losses in low current areas versus persistence in recent years in intermediate and high current areas. Areas of intense mixing may constitute kelp refugia from physical stressors, and low current areas may exacerbate the negative effects of stressors.

Currents can also mediate biotic stressors. In the San Juan Archipelago in northern Puget Sound, mesograzers, especially the small snail *Lacuna vincta,* play an important role in mortality to *Nereocystis* in hydrodynamically quiescent habitats; periods of weak currents allow grazers to crawl up and structurally damage stipes, making them vulnerable to shear under strong, infrequent tidally driven drag force [34]. While *Lacuna* snails were not commonly observed in SPS in 2017 and 2018, kelp crabs (*Pugettia producta*) were abundant on the blades, bulbs and stipes in the *Nereocystis* forests that were not subjected to regular, intense currents. Kelp crabs preferentially consume fresh *Nereocystis* in Puget Sound, and laboratory and field experiments suggest that they may play an important role in mediating the growth and survival of *Nereocystis* in the Salish Sea [139, 140]. Sea urchins can control kelp populations, especially in the absence of predators, however sea urchins were observed to be absent or rare in SPS.

Many other factors that are known to drive kelp abundance, and are outside the scope of this study, likely also played a role in the observed changes in *Nereocystis* distribution. Human activities—especially logging and coastal development—have increased sediment [141–144], nutrient[135] and pollutant loads to coastal ecosystems [134]. These factors are associated with the global ‘flattening of kelp forests’, through altering competitive interactions with turf algae [20]. In SPS, widespread deforestation began in the mid-1850s, and likely profoundly increased sedimentation. Changes to nearshore biotic interactions, often through fishing/harvest, can alter controls on grazer populations by decreasing predation [6, 17, 18]. In SPS, rockfish [145] and other groundfish [146], salmonids [147] and forage fish [148] populations have been dramatically reduced relative to historical levels. These species occupy middle to high trophic positions, directly and indirectly influencing populations of kelp grazers [18, 149]. Alterations to disturbance regimes following changes in trophic dynamics can also facilitate competition between *Nereocystis* and other macroalgal species. In the absence of disturbance, perennial algae can exclude annual kelp species such as *Nereocystis* [17]. The invasive perennial alga *Sargassum muticum*, which was observed at many historical and current *Nereocystis* sites in SPS, can competitively exclude native kelp through shading [25]. Compounding the effects of these diverse stressors, sporophyte mortality may impact basin wide bed connectivity because most spores settle within a few meters of the parent sporophyte [150].

While the historical data in this study lacked sufficient temporal resolution for comparison to climate indices, two notable climate events corresponded to observed *Nereocystis* declines. The large decline following the 1978 survey coincided with a shift from a cold to a warm regime defined in Pacific Decadal Oscillation (PDO) data [151]. The decline between 2013 and 2018 coincided with a period of warm sea surface temperature in the northeast Pacific known as ‘the Blob’ [152]. Marine heatwaves doubled globally between 1982 and 2016 and are projected to become more frequent and extreme [153].

## Conclusion

This study identified substantial losses of *Nereocystis*, an important biogenic habitat, in SPS over 145 years. It also related several physical factors that are commonly associated with kelp performance to the temporal variability in *Nereocystis* extent. Given the importance of biogenic habitats to ecosystem structure, the findings underscore a need to investigate the potential causes of change, possible management responses, and linkages to other species.

## Acknowledgments

This work was funded by the Washington State Department of Natural Resources, Washington Coastal Resilience Project, and U.S. Geological Survey Coastal Habitats in Puget Sound Project. We thank many scientists who contributed to field work and data synthesis, including Jeff Gaeckle, Julia Ledbetter, Kate Sherman, Olivia Hannah, Melissa Sanchez, and Lauren Johnson. We thank Tarang Khangaonkar, Christopher Krembs and Julia Bos for providing environmental data and manuscript review. We thank Cathy Pfister and Megan Dethier for insightful comments on earlier versions of this manuscript. HB designed the study, collected and analyzed the data, and wrote the paper with input from all authors. TM, BC, MC, LF, EG and NV collected and analyzed data and contributed to writing. PD assisted with analysis and writing.

## Supporting Information

**S1 File. Data used in this study.** Excel spreadsheet with data used in all analyses.

**S1 Text Comparison of synoptic survey methods and results.**

**S1 Fig. Distribution of the length of individual *Nereocystis* features in comprehensive snapshot surveys.** The distribution of kelp bed feature length in six comprehensive surveys, ranging from a median of less than 0.1 km (2017) to 1.5 km (1911). Differences in length are likely to be related to both survey resolution and actual length of kelp features.

**S2 Fig. Linear extent of shoreline with *Nereocystis* present between 1876 and 2017, based on six comprehensive snapshot surveys, summarized over three sub-basins.** Recent estimates (1998 and 2017) are dramatically reduced relative to estimates in 1876, 1935 and 1978. The 1911 estimate could represent a low point in kelp extent, but likely reflects methodological differences in survey methods. This depiction of linear extent shows similar results as the number of 1-km segments with *Nereocystis* present (Fig 4).

**S3 Fig. Nearshore temperature profiles on July 18, 2018 at seven sites in SPS.** Casts were collected at approximately – 6 m (MLLW). At Squaxin Island, Devil’s Head, Day Island and Salmon Beach, three casts were collected, at stations in the center of the site and near the alongshore site boundaries. At other sites, single casts were collected at the center of the site. During this summer period of maximum expected stratification, the profiles showed little change in temperature with depth. Multiple casts at up to three stations per site, showed similar values within site and a gradual gradient among sites.

